# Quantifying polygenic effects in genome-wide association studies using generalized estimating equations

**DOI:** 10.1101/032854

**Authors:** Julian Hecker, Dmitry Prokopenko, Christoph Lange, Heide Löhlein Fier

## Abstract

Recently, LD Score regression^1^ has been proposed as a computationally fast method to contrast confounding biases with polygenicity and to quantify their contribution to the inflation of test statistics in GWAS.

In this communication, we extend the LD Score regression approach by applying the generalized estimation equations (GEE) framework, which is capable of incorporating more external information from reference panels about the correlation structure of test statistics. We apply our GEE approach and LD Score regression to simulated and real data to compare their performance.

We show that our proposed methodology obtains more efficient estimates while preserving the robustness and desired properties of LD Score regression.

## 2 Introduction

In their seminal paper, Yang et al. (2011) have revealed that a substantial inflation of test statistics in genome-wide association studies can be attributed to the presence of polygenic inheritance ^2^.

For a specific variant, polygenic inheritance inflates the test statistic proportional to the amount of genetic variation of a disease susceptibility locus (DSL) that the variant tags.

Recently, Bulik-Sullivan et al. (2015) have shown that the effect of confounding biases on the inflation of test statistics does not correlate with the genetic variation that the variant captures^1^. Confounding biases refer to the presence of technical artefacts as cryptic relatedness and/or population stratification.

This observation has been the motivation for so-called LD Scores, estimated from the 1000 Genomes reference (1kG) panel^3^, in order to derive their mean model for the test statistic of variant *j* in a quantitative trait study:

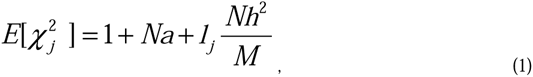

where *N* is the sample size, 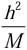 the average heritability per SNP, *M* the number of SNPs and *1_j_* is the LD Score. The parameter *a* measures confounding biases. With minor modifications, this model also holds for case-control-studies (see supplementary material^1^).

To estimate the parameters of interest, the method of Bulik-Sullivan et al. (2015) uses a weighted linear regression, restricted to common, and usually well imputed variants. The weights for the regression are introduced to reduce the standard error by correcting for correlated test statistics and heteroskedasticity. Standard errors for the LD Score regression estimates are obtained by a block jackknife method. Under some assumptions about the effect sizes, it is possible to derive an estimate of the heritability from the polygenic term^1^.

Our approach builds on the same mean model as the LD Score regression, but applies a different estimation procedure, which incorporates more external information about the correlation between test statistics. Our proposed methodology therefore achieves higher efficiency and can calculate valid standard errors without bootstrap methods.

In order to analyze summary statistics from association studies with large sample sizes, several publications use the multivariate normal distribution framework of z-scores (Connelly and Boehnke, 2007^4^; Han et al., 2009^5^; Wen and Stephens, 2010^6^; Zaitlen et al., 2010^7^).

Motivated by this framework, the correlation between two specific *χ*^2^ test statistics of two variants can be derived as the LD measure *r*^2^ between both loci.

As exploited by Bulik-Sullivan et al. (2015) for the calculation of the LD Scores, the majority of LD vanishes after a genetic distance of approximately 1 centi morgan (cM) (supplementary table 10 in Bulik-Sullivan et al. 2015)

Combining both observations, we can conclude that only spatially close test statistics are correlated.

The LD measure *r*^2^ can be estimated from a reference panel. But due to relatively small sample sizes of reference panels, the correlation matrix for multiple test statistics can only be estimated accurately for a moderate-sized group of test statistics (Wen and Stephens, 2010).

Since LD differs between populations, the LD structure from a reference panel only approximates the correlation structure between statistics in a real study. Additionally, the incorporation of confounding covariates into the association analysis can also affect the correlation structure between statistics^8^.

Motivated by these observations, we apply and extend the framework of generalized estimating equations (GEEs) to estimate the parameters of the mean model and obtain asymptotic valid standard errors. In more detail, we split the human genome into blocks of 1cM length and group test statistics within these blocks into clusters. This block strategy respectively the estimation of block correlation matrices, is also applied for the imputation of summary statistics (Pasaniuc et al., 2014^9^; Lee et al., 2013^10^), or the gene-based test VEGAS^11^.

We then conclude that only spatially close clusters are correlated and that the correlation structure within a cluster can be well approximated by the LD matrix estimated from the 1kG reference panel.

Using these LD matrices as the working-correlation matrices for each corresponding cluster in the GEEs, we achieve higher efficiency compared to LD Score regression. In the supplementary material we show that we can overcome the violated GEE-assumption of independent clusters and are able to extend the sandwich-covariance estimator to this scenario of sparsely correlated clusters.

It is important to emphasize, that the LD matrices do not perfectly have to match the true correlation structure of test statistics in order to obtain consistent estimates and asymptotic valid standard errors.

Analogously to the LD Score regression, it is also possible for our method to constrain the intercept in (1) to a specific value and to only estimate the polygenicity term.

For further details of the derivation of our methodology, we refer to the Methods section and the supplementary material.

## 3 Methods

### LD Scores and input

As described in the introduction, our approach is based on the same mean model as the LD Score regression (see Equation (1)^1^) Therefore, we adopt the LD Scores as they were defined in Bulik-Sullivan et al. (2015). The LD Scores for the European-ancestry samples in the 1kG project are available from the LD Score regression Web page. For the details of the calculation, we refer to these Web resources.

We suggest to restrict the input of test statistics to variants that are a subset of the HapMap3 SNPs^12^ variants are usually well imputed. If imputation info-scores are available, we filtered with info-score > 0.9.

### Exponential family modeling of test statistics

As explained in the introduction, our approach is based on the generalized estimating equations (GEE) framework. It is natural to derive the GEE-related objects under the assumption that the distribution of the test statistics is described by an exponential family distribution.

We combine the mean model in equation (1) with the additional assumption that the variance of a test statistic is given by

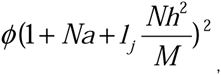

where *ϕ* is a overdispersion parameter.

This formulation is motivated by the assumption of normally distributed effect sizes, as used in the derivation of the linear mixed model of GCTA^13^. This would imply *ϕ =* 2. The heteroskedasticity weights of the LD Score regression were also motivated by this assumption.

This leads in our scenario to the description of test statistics distribution via the gamma distribution. Note that we only use assumptions about the moments, not the distribution in general.

### GEE objects and asymptotic results

In the supplementary material, we derive the corresponding objects to set up the GEEs. These equations are used to estimate the intercept and the polygenicity term. See below for details of the Implementation.

In addition, in the supplementary material we state the technical assumptions under which we can establish asymptotic results for the estimation. We allow the number of parameters to grow with the number of blocks with some specified speed. The critical assumption is that the blocks are only sparsely correlated, that means only spatially close blocks. More precise, we assume that the number of correlated blocks is *O(n)*. An important result is the derivation of a consistent covariance estimator via a modified sandwich estimation. This estimator is used to obtain valid asymptotic standard errors.

### Implementation and running time

Map information about the genetic distances are available through the 1kG data and are included in the LD Score files. After the determination of the blocks along the chromosomes and the filtering of input test statistics, we estimated the working correlation matrices from 1kG haplotype data for the European-ancestry samples. We ensured that these matrices were positive definite by slightly scaling them towards the identity matrix. After setting up the related objects, the GEEs were solved via the Fisher score algorithm. The covariance estimation was then calculated with the corresponding expression in the supplementary material.

By extracting LD information from chromosome to chromosome, the memory requirements are below 3gb and the running time for estimation of parameters and standard errors takes less than 3 minutes.

## 4 Results

### Robustness of the mean model and LD Scores

Bulik-Sullivan et al. (2015) investigated the stability of the LD Scores across the European-ancestry populations and the behavior of the LD Score regression in simulation scenarios with confounding biases, polygenicity or both. The LD Score regression performs robustly in all these scenarios.

Since these properties are only related to the LD Scores and the mean model (1), we conclude that their results transfer to our approach. We refer to the results section in Bulik-Sullivan et al. (2015) for more details.

### Simulation study

To set up a realistic simulation scenario, we used LD Scores, calculated by the ldsc software (Bulik-Sullivan et al., 2015), over the European subsample of the 1kG project. These LD Scores measure the genetic variation tagged by the corresponding variants.

On chromosome 2, we identified about 100,000 variants, that are included in the HapMap 3 project^12^, the LD Score data set and with haplotype data from the 1kG project available.

We randomly selected 62,500 variants as a reasonable coverage of filtered, well imputed data.

The genetic positions along these variants ranged from approximately 0 to 275 cM.

Due to computational restrains while handling large matrices, we partitioned the chromosome in 11 regions of about 25 cM length.

For each region, we estimated the LD matrix between the variants from the haplotypes and truncated LD to zero between variants with more than 1 cM distance.

We used the resulting LD matrix as the correlation matrix of normally distributed z-scores with variance

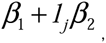

where *1_j_* is the LD score for the corresponding variant. We assigned multiple combinations of reasonable values for the two parameters, in order to simulate a real data set.

To achieve the setting of genome-wide data, we repeated 13 draws of these variants and obtained 812,500 z-scores resp. squared test statistics.

We simulated 1,000 replications to compare our approach with the LD Score regression.

The mean model was determined by equation (1) and is the same for both methods.

We implemented a weighted linear regression as used by the LD Score regression. Recall that the weights of the LD Score regression are given by (see Online Methods in Bulik-Sullivan et al. (2015)):

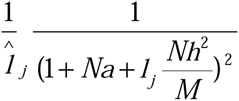

The overcounting weights 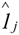 are the LD Scores, calculated only over regression SNPs. We used the LD matrices for each 25cM region above to calculate these weights for each variant. To simplify, we used the true parameters for the heteroskedasticity weights of the LD Score regression. In practice, these are estimated by the LD Score regression software in a first step.

For the GEE approach, we split each region into 1cM blocks, resulting in about 25 blocks for each region. For each block, we estimated the correlation structure between the test statistics from the LD matrix for this region.

In table I we listed the results from the estimation over these 1,000 replications for 8 combinations of parameter values. Both methods estimated the parameters consistently, we only report variances resp. standard errors.

**Table I:**
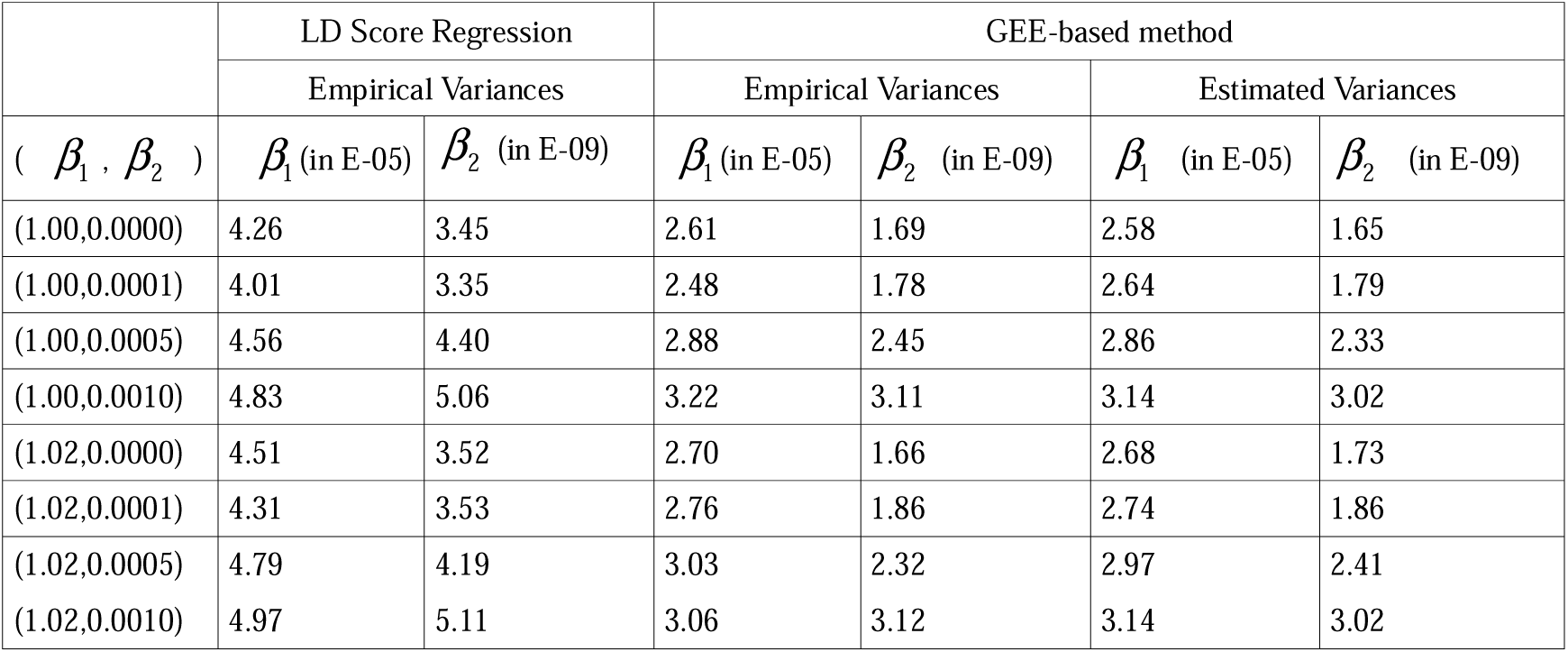
Comparison of both methods in a simulation study for 8 different parameter configurations. In this table we listed the empirical variances over the 1,000 replications resp. the GEE-sandwich estimated variances of the parameter estimates.

|  | LD Score Regression |  | GEE-based method |  |  |  |
| --- | --- | --- | --- | --- | --- | --- |
|  | Empirical Variances |  | Empirical Variances |  | Estimated Variances |  |
| ( $\beta_1, \beta_2$ ) | $\beta_1$ (in E-05) | $\beta_2$ (in E-09) | $\beta_1$ (in E-05) | $\beta_2$ (in E-09) | $\beta_1$ (in E-05) | $\beta_2$ (in E-09) |
| (1.00,0.0000) | 4.26 | 3.45 | 2.61 | 1.69 | 2.58 | 1.65 |
| (1.00,0.0001) | 4.01 | 3.35 | 2.48 | 1.78 | 2.64 | 1.79 |
| (1.00,0.0005) | 4.56 | 4.40 | 2.88 | 2.45 | 2.86 | 2.33 |
| (1.00,0.0010) | 4.83 | 5.06 | 3.22 | 3.11 | 3.14 | 3.02 |
| (1.02,0.0000) | 4.51 | 3.52 | 2.70 | 1.66 | 2.68 | 1.73 |
| (1.02,0.0001) | 4.31 | 3.53 | 2.76 | 1.86 | 2.74 | 1.86 |
| (1.02,0.0005) | 4.79 | 4.19 | 3.03 | 2.32 | 2.97 | 2.41 |
| (1.02,0.0010) | 4.97 | 5.11 | 3.06 | 3.12 | 3.14 | 3.02 |

The entries in the first 4 columns of the table give the empirical variances for the corresponding method. In the last two columns we stated the estimated estimation variance for the GEE approach, calculated by our sandwich-covariance formula.

First, we conclude that that our covariance estimates correctly estimates the variances of our GEE method.

Second, we see that the LD Score regression estimation variance is up to factor 2 larger than the estimation variance of our GEE-based approach. To emphasize the advantage of this variance reduction, we considered the parameter configuration with small polygenic effect *β*_1_ = 1.0, *β*_2_ = 0.0001. If we use the estimate of *β*_2_ and the corresponding estimated standard error, both from our GEE-based method, to test the hypothesis that the polygenic effect is 0, with reference to the significance levels *α =* 0.05,0.01,0.001, we observe a estimated power, based on the 1,000 replications, of 67.4%, 42.6% resp. 16.9%. If we use the estimate of *β*_2_ from the LD Score regression and correct with the corresponding estimated empirical standard error over the 1,000 replications, we obtain a power of only 41.0%, 20.2% resp. 6.1%.

### Real data

In order to compare the performance of our method in relation to the LD Score regression, we analyzed the public available summary statistics from the Psychiatric Genomics Consortium (PGC). In particular, we considered the data sets for the five psychiatric disorders Bipolar Disorder (BIP)^14^, Schizophrenia (SCZ)^15^, Major Depressive Disorder (MDD)^16^, Attention Deficit Disorder (ADHD)^17^ and Autism Spectrum Disorder (AUT) from the Cross-Disorder Group^18^.

First, we filtered the summary statistics with the ldsc software, using the default parameters^1^. This included removing ambiguous SNPs and variants with an imputation info-score < 0.9.

Afterwards, we removed variants that are not included in the ref-LD-Score set resp. weight-LD-Score set, not part of the HapMap3 SNPs^12^ or without haplotype information from the 1kG-phase1 dataset. The final numbers of variants for the analysis are listed in table II. This procedure ensured that we only considered well imputed variants and could use the same input for both methods.

**Table II:** Comparison of both methods based on the analysis of real datasets. We listed the estimated values for both parameters and the method-based estimated variance in brackets.

|  |  | LD Score regression |  | GEE-based method |  |
| --- | --- | --- | --- | --- | --- |
| Disorder | #SNPs | 1+Na | Polygenic term | 1+Na | Polygenic term |
| BIP | 812,596 | 1.0318(7.57E-05) | 0.000597(5.70E-09) | 1.0323(5.47E-05) | 0.000534(4.49E-09) |
| SCZ | 840,481 | 1.0418(1.37E-04) | 0.001091(1.40E-08) | 1.0445(8.02E-05) | 0.000954(9.58E-09) |
| MDD | 888,591 | 1.0184(6.89E-05) | 0.000311(5.12E-09) | 1.0167(4.21E-05) | 0.000278(3.49E-09) |
| ADHD | 803,096 | 1.0010(6.72E-05) | 0.000056(4.82E-09) | 0.9973(4.31E-05) | 0.000063(2.77E-09) |
| AUT | 936,044 | 0.9898(5.63E-05) | 0.000451(4.16E-09) | 0.9888(4.08E-05) | 0.000412(3.86E-09) |

In table II, we present the estimated parameters for the intercept and the polygenic term for both methods. In brackets, the estimated variance is listed. The estimated variance of the LD Score regression was calculated using the default parameters of the ldsc-software. We can observe that the variance of the parameter estimates for the LD Score regression is up to 75% larger.

### Discussion

Based on the proposed mean model by Bulik-Sullivan et al. (2015), we introduce a more efficient GEE-based framework to estimate the contributions of confounding biases and polygenicity to the inflation of test statistics in GWAS. The increased efficiency of our approach is achieved by incorporating local LD information from an external reference panel, while our GEE approach does not require that the reference panel and the study data have exactly the same LD structure. It is robust against deviations of the sample LD structure from the reference panel. Since we use the same mean model as Bulik-Sullivan et al. (2015), our method preserves the desired properties of the LD Score regression. As described in^1^, the estimated intercept of the mean model provides a more robust quantification of the extent of confounding biases than the Genomic Control *λ_GC_*_19_. In conclusion, our approach improves the estimation framework of the LD Score regression method with reasonable additional computational effort.

Our theoretical derivations and assumptions are quite general. In particular, we allowed the number of parameters to grow with the number of blocks. For further research, this makes it possible to extend the mean model to incorporate more heritability components or estimate the genetic correlation between two traits.

Another interesting point is the possible incorporation of the results in Xu et al.^8^ to obtain a better approximation of the correlation structure.

